# Fc-independent SARS-CoV-2 infection-enhancing antibodies decouple N-terminal and receptor-binding domains by cross-linking neighboring spikes

**DOI:** 10.1101/2024.08.21.608921

**Authors:** Floris J. van Eerden, Songling Li, Tina Lusiany, Hendra S. Ismanto, Tohru Terada, Christoph Gerle, Kanako Akamatsu, Mika Hirose, Fuminori Sugihara, David Virya Chen, Jun-ichi Kishikawa, Takayuki Kato, Yafei Liu, Masato Okada, Hisashi Arase, Daron M. Standley

## Abstract

Antibody dependent enhancement (ADE) is a serious concern in vaccine development. The canonical ADE pathways are dependent on the fragment crystallizable (Fc) region of the antibody. In SARS-CoV-2 several antibodies have been discovered that inflict ADE in vitro. These antibodies target the N-terminal domain (NTD) of the SARS-CoV-2 spike protein. We previously proposed that these NTD-targeting infection-enhancing antibodies (NIEAs) cross-link neighboring spike proteins via their NTDs, and that this results in a decoupling between the NTD and receptor binding domain (RBD), facilitating the “RBD down” to “up” transition. In this study we present a combination of molecular dynamics simulations and cryogenic electron microscopy data that, together, demonstrate that NIEAs are indeed able to cross-link neighboring SARS-CoV-2 spike proteins, and that this cross-linking results in a decoupling of the NTD and RBD domains. These findings provide support for an Fc region independent ADE pathway that is not only relevant for SARS-CoV-2 but also for other viruses of which the spike proteins undergo a conformational change upon host cell entry.

## Introduction

The SARS-CoV-2 virus has taken a significant toll on global health and economy. In spite of the remarkable success of vaccine development, the virus continues to evolve and people continue to be infected, with many COVID-19 patients experience long-term symptoms^1–3^. The severity of COVID-19 has been shown to depend on adaptive immune responses, particularly those mediated by B cell lymphocytes^4,5^. Consistently, not all B cells produce neutralizing antibodies^6–11^; indeed, a distinct subset of anti-SARS-CoV-2 antibodies have been shown, instead, to enhance infection^12,13^. Antibody dependent enhancement (ADE) has been observed in a number of diseases and is an important consideration in vaccine design^14–21^. The canonical ADE pathway involves host cell entry via fragment crystallizable (Fc) receptors on the surface of the cell^14^. While several other ADE pathways have been described^22–25^; all those reported prior to the COVID-19 pandemic depend on the Fc region of the antibody.

In the case of SARS-CoV-2, two types of infectivity enhancing antibodies have been identified: one targeting the RBD region that is Fc-region dependent; in addition, a second type of antibody that targets the N-terminal domain (NTD) of the spike protein has been observed^12,13^. These NTD-targeting infection-enhancing antibodies (NIEAs) constitute a distinct subtype that is Fc region independent. NIEAs have been shown to promote the transition of the RBD from the “down” to the “up” orientation, which exposes the ACE2 binding interface, which is a prerequisite for RBD-ACE2 binding^8,10^, although the mechanism of this effect remains obscure. Since the NTD domain interacts with the RBD in its down conformation^8,11^, a local displacement of the NTD away from the RBD may facilitate the RBD-down to up transition^26–30^.

We previously proposed a model in which cross-linking the NTDs of two neighboring spikes by a single NIEA would exert an outward force on the NTDs, resulting in NTD-RBD decoupling^13,31^. In short, this model is supported by several observations. First, while full IgG or diFab fragments of NIEAs enhanced ACE2 binding by SARS-CoV-2^13^, a single Fab fragment has no effect^13^, which suggests that both NIEA Fab arms have to be NTD-bound in order to enhance SARS-CoV-2 infectivity. These observations also suggest that the Fc region is dispensable, which has indeed been demonstrated^13^. Structural modelling further showed that it is unlikely that a single NIEA can bind two NTDs on a single SARS-CoV-2 spike at the same time^13,31^. Finally, the spike-spike spacing observed in EM tomography^32^ agrees well with a static model of two NIEA-cross-linked model spikes, for which the spacing is 23 nm. Taken together, the cross-linking model is consistent with a wide range of independent observations. Nevertheless, NIEA cross-linked spikes have not been observed directly.

In this study we used a combination of electron microscopy and molecular dynamics to investigate NIEA cross-linking and its influence on the strength of RBD-NTD interactions. As a preliminary step in this aim, we recently reported negative staining electron microscopy images of NTDs bound by the two arms of a single NIEA that agree with our proposed cross-linking model^31^. Here, we have extended this EM study using full spike trimers in order to observe a pair of cross-linked spikes for the first time. We then carried out extended MD simulations in order to observe the effect that cross-linking has on NTD-RBD interaction as a function of time. By using a combined experimental-computational approach we could construct a realistic picture of how NIEA cross-linking decouples the NTD from the RBD, thereby facilitating SARS-CoV-2 infection in an Fc region independent manner.

## Results

### Electron microscopy shows that NIEAs cross-link soluble SARS-CoV-2 spikes

We obtained strong support for the notion that NIEAs can cross-link SARS-CoV-2 spike proteins using negative staining electron microscopy. While spike trimers were dispersed in solution, addition of equimolar NIEA caused spikes to cluster in rosettes (Figure 1A). The appearance of rosettes upon addition of NIEA suggest the consecutive cross-linking of spike trimers joined at the base of their stalks.

**Figure 1:**
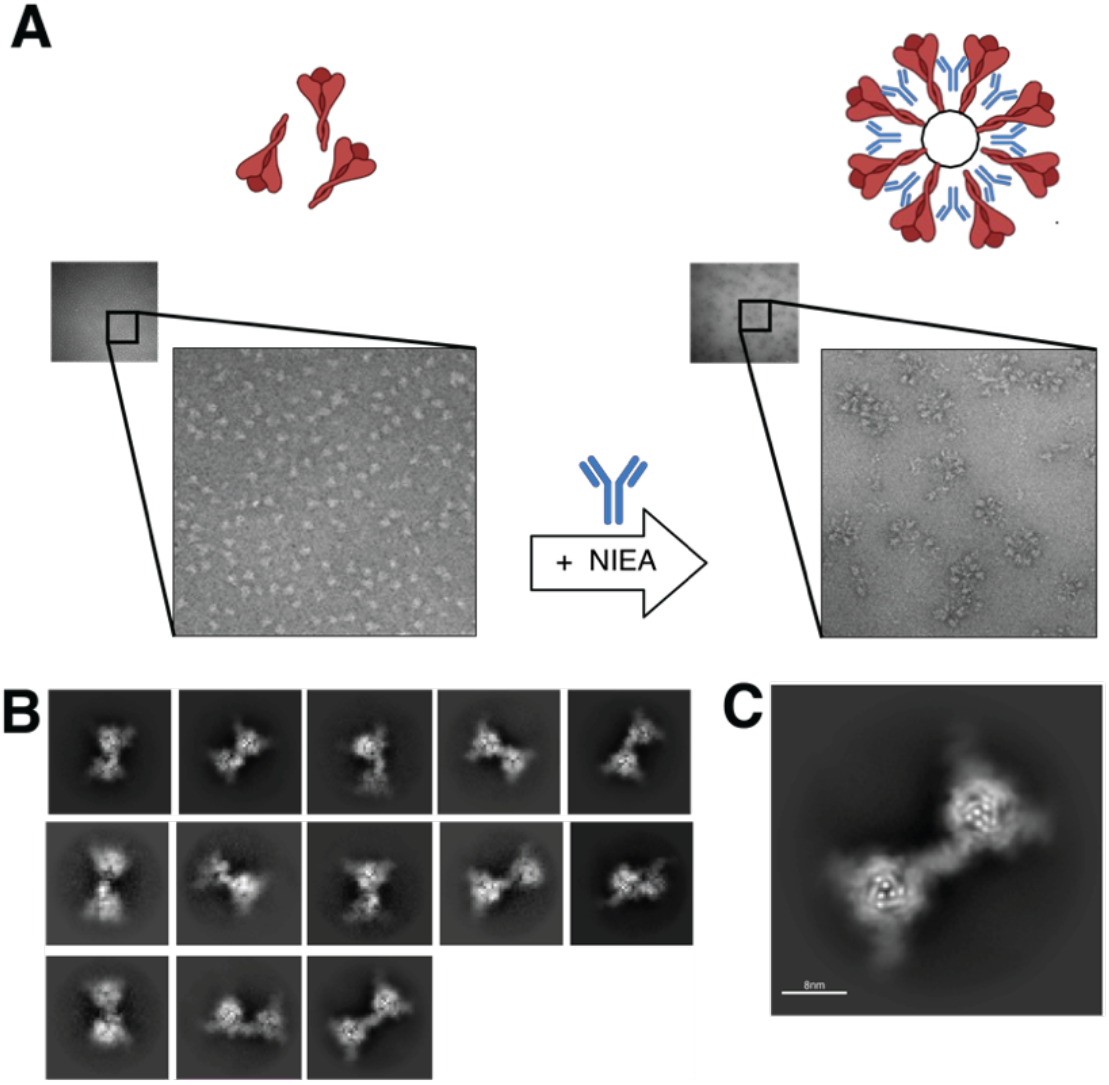
Electron microscopy imaging of SARS-CoV-2 spikes cross-linked by NIEA. (A) Addition of NIEA to SARS-CoV-2 spike protein in solution results in the cross-linking of the spike proteins, illustrated in upper panel, and observed by negative staining electron microscopy in the lower panel. Schematic diagram was created with BioRender.com. (B) Representatives of 2D classes obtained after particle re-picking. (C) Best 2D class depicting trimeric spike cross-linked by full length NIEA

We next observed the cross-linked complexes by cryogenic EM (cryo-EM). To avoid rosette-like complex formation, we reduced the spike:NIEA ratio from equimolar to 2:1. During the initial image analysis we observed density maps and 2D classes of NIEA Fabs bound to spikes (supplemental Figure S1A). In order to visualize the IgG density in these 2D class images, we next re-extracted the particles and shifted the origin of the particle box from the center of the spike trimer to the center of NIEA (supplemental Figure S1B). In this manner we obtained 2D classes in which two spike proteins were cross-linked by a single NIEA (Figure 1B). One class in particular clearly showed the cross-linking of two spike trimers by a single NIEA (Figure 1C). We estimated the distance between spike centers to be approximately13.9 nm. This image is the first direct observation of SARS-CoV-2 spike cross-linking by NIEAs.

### Modelling of the SARS-CoV-2 NIEA cross-linked spike system

To understand the structure and dynamics of the NIEA-cross-linked spikes, we next prepared an atomistic model of two SARS-CoV-2 spike proteins cross-linked by an NIEA. The system was prepared in three steps. First, all-atom modelling of an NTD bound by a NIEA Fab fragment was carried out. In this step, the previously-reported NTD-Fab structure was utilized^13^, glycans were added and clashes were removed using a rotamer library^33^. Second, the NTD was extended to a full-length spike trimer. In this step, the SARS-CoV-2 spike model from Casalino and co-workers^34^ was used. One of the three NTD domains was selected at random and replaced by the NTD-Fab from the first step. Subsequently, the actual cross-linked system was built from two Fab bound SARS-CoV-2 spikes, where special care was taken to create a system in which the NIEA was in a natural position. Two copies of the Fab bound SARS-CoV-2 spike were positioned on a plane (Figure 2A). For each spike the principal axes were determined, along with another axis between the C-termini of the heavy chains of the Fab fragments. Subsequently the positioning of the spikes was established by determining their relative rotation as well as the distance between the spikes. The spikes were rotated stepwise along the longest principal axis (Figure 2B) and the distance between the spikes was sampled along the axis connecting the Fab C-termini (Figure 2C). For each combination of rotation and distance, the Fc region of the NIEA was reconstructed^35^, and the models were ranked according the RMSD of the two Fab fragments after reintroduction of the Fc region. The model with the lowest Fab RMSD after remodeling the antibody constant domain was selected as a starting point for subsequent molecular dynamics simulations. The distance between the two spikes, as measured by the Proline 499 residues in the NIEA bound chains, was 21.7 nm, very close to that observed in EM tomography^32^. These meticulous model building steps insured that the entire system was in a relaxed structural arrangement.

**Figure 2:**
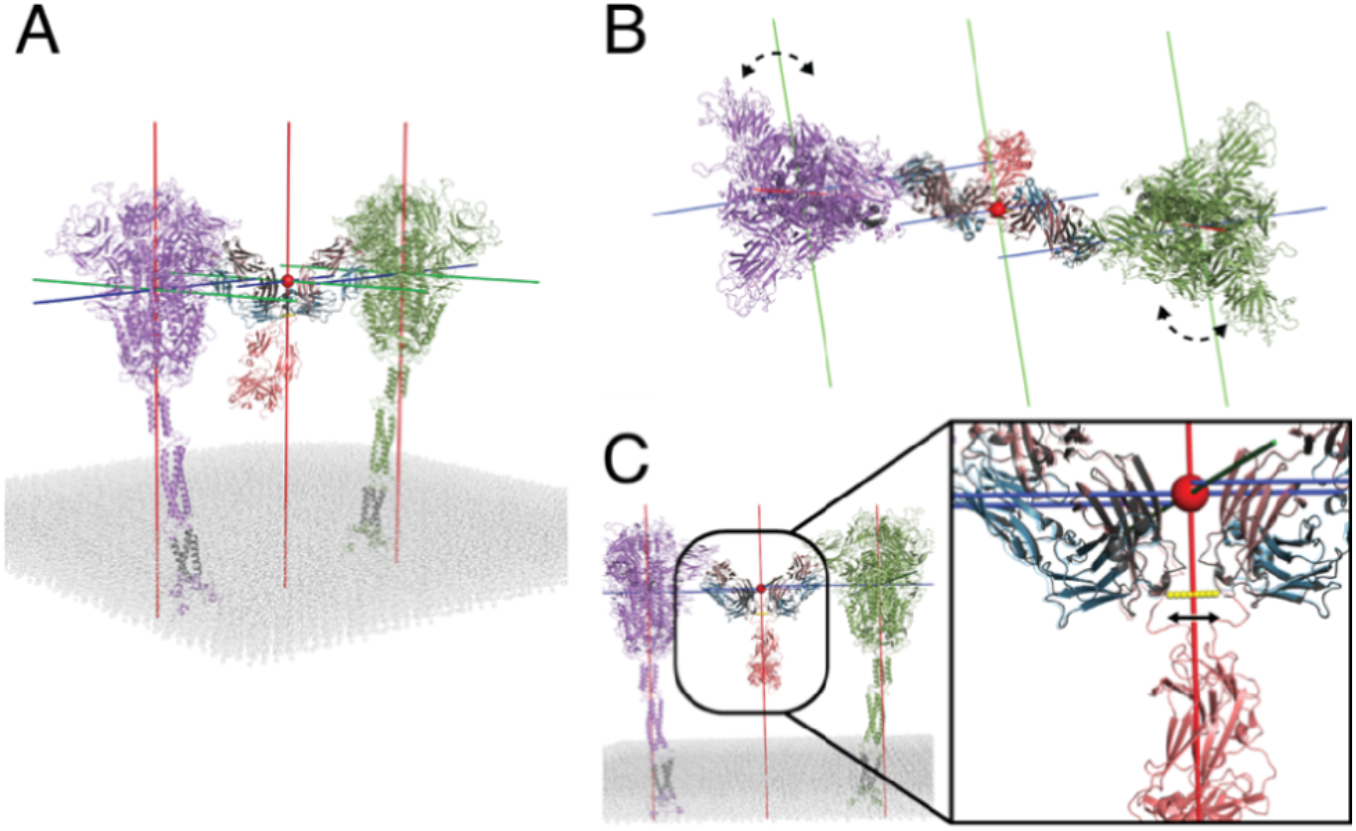
Modelling of SARS-CoV-2 spikes cross-linked by NIEA. Care was taken to create a system in a realistic configuration. (A) Two SARS-CoV-2 spike with Fab fragment embedded in a planar membrane slap, with reconstructed Fc region. Principal axes of the spikes and NIEA are indicated. (B) Spikes were rotated along their longest principal axis, to sample their relative rotational sampling. (C) Distances between the two spikes was varied along the axis connecting the heavy chain C-termini of the Fab fragments, see inset.

### Overview of simulation systems

The obtained model of the cross-liked NIEA spike system was subsequently prepared for molecular dynamics simulations; the bilayer slab was replaced by a membrane mimicking the endoplasmic reticulum-Golgi intermediate compartment, which is the membrane from which the virus is released^13,34,34^, and both the spikes and the NIEA were decorated with glycans (Figure 3A-B). The primary interface between NTD and RBD domains is through neighboring chains: NTD chain A – RBD chain C, NTD B – RBD A, NTD C – RBD B (Figure 3C,D). In our system, the A-C pair was NIEA-bound. We thus hypothesized that spike cross-linking via a NIEA results in a de-coupling of the NTD A-RBD C interaction, compared to the other NTD-RBD pairs.

**Figure 3:**
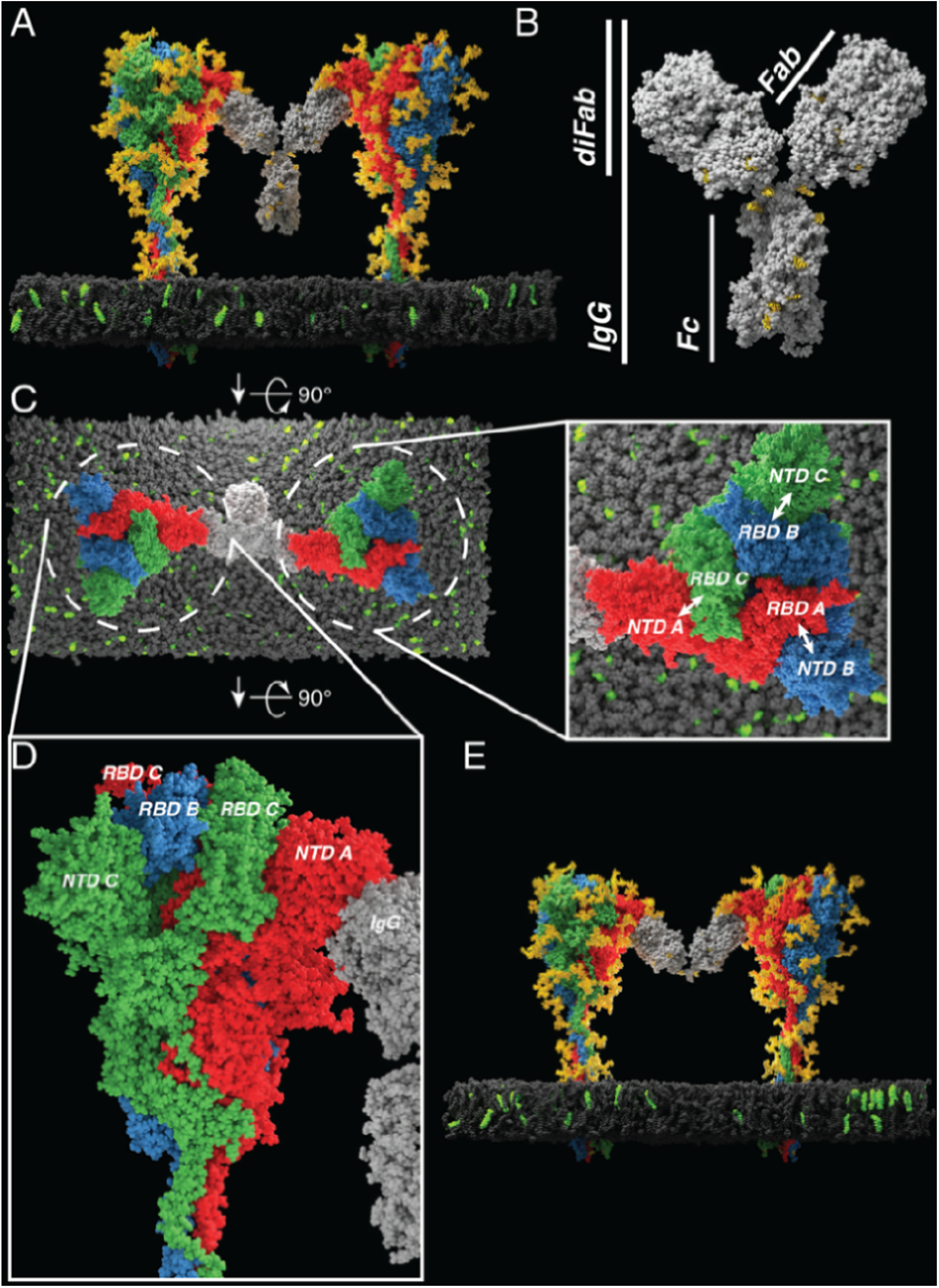
Cross-linking model infectivity enhancing antibodies. (A) Simulation systems consisting out of two SARS-CoV-2 spike proteins bound by an IgG. Spikes are colored by chain, glycans are rendered in yellow. Spikes are embedded in a membrane (grey) with a composition mimicking the viral membrane, cholesterol is shown in bright green. IgG is colored in light grey. Solvent is not shown for visibility. The left spike corresponds to spike 1 and the right one to spike 2 in the RMSD analysis. (B) Rendering of the COV-2490 IgG with the different sections of the antibody indicated: IgG, Fab, diFab, and the Fc region. Protein is rendered in grey with glycans in yellow. (C) Top view of the IgG simulation system, with the inset indicating the NTD and RBD domains. Glycans and solvent are not shown for visibility. (D) Side view of the crown of the spike protein, illustrating how the NTD and RBD domains interlock. Glycans and solvent not shown for visibility. (E) Simulation system of two SARS-CoV-2 spike proteins bound by a diFab, coloring as in panel A.

To investigate the role of the Fc region, a second system was constructed in which the spike proteins were cross-linked by a diFab instead of a full IgG (Figure 3E). For both the IgG and diFab systems, ten independent simulations were performed, with simulation times ranging from 357 ns to 1.1 μs. RMSD traces of the spikes showed that there was little variation between the three polypeptide chains in the simulation, indicating that the effect of the IgG binding is subtle in comparison with the overall motion of the spike (Figure 4A).

**Figure 4:**
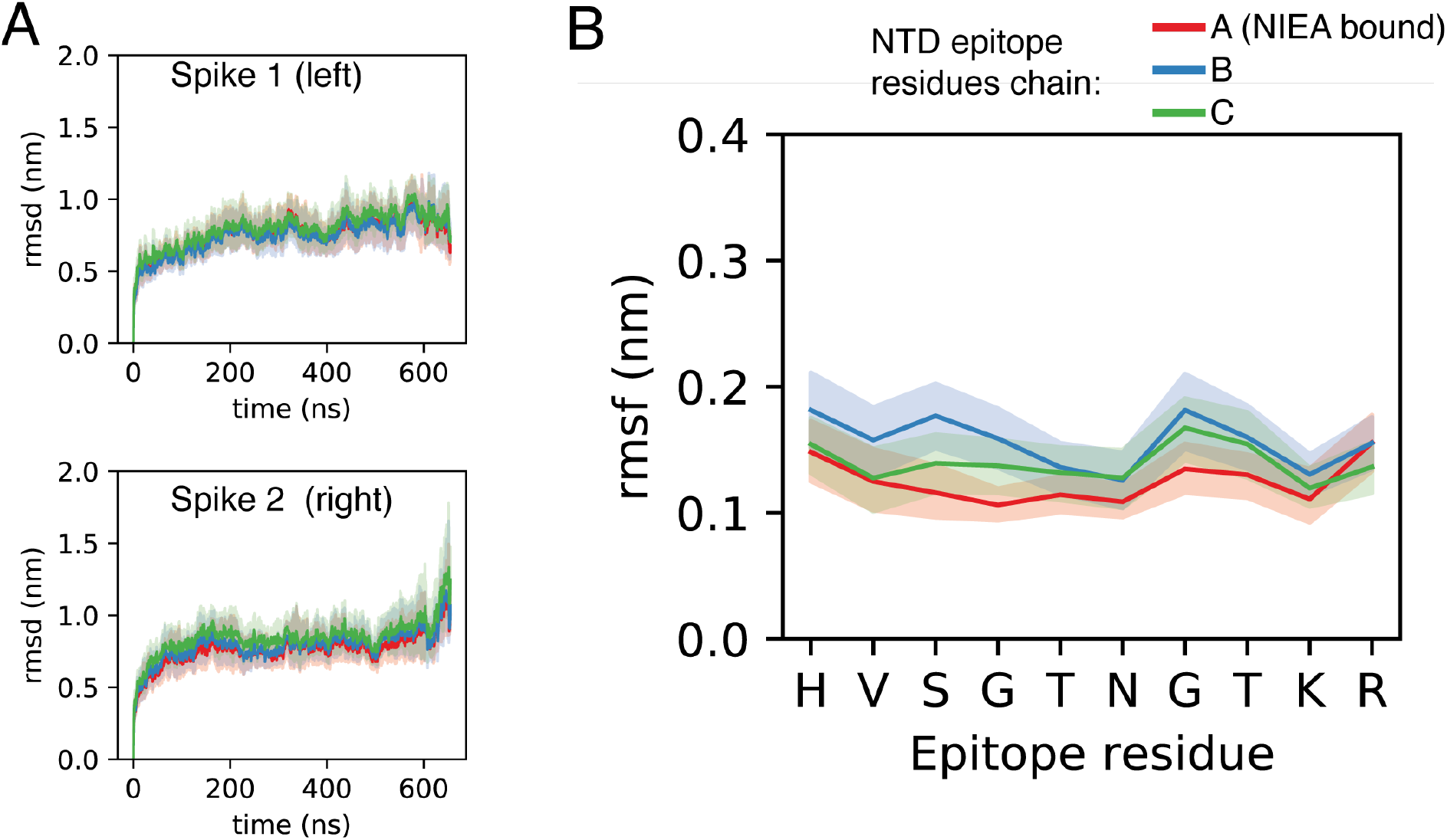
RMSD and epitope RMSF of the SARS-CoV-2 spike protein in the molecular dynamics simulation. (A) Per chain RMSD of the SARS-CoV-2 spikes. The red trace represents the NIEA bound chains. The per chain RMSD was calculated by fitting the trajectory on the whole spike protein. Mean and 95% confidence interval are shown. Spike 1 (left) corresponds to the left spike in Figure 3A, and Spike 2 to the right one. Chain A (red), chain B (blue) and chain C (green). Confidence interval was calculated using Seaborn ^36^. (B) RMSF of the SARS-CoV-2 epitope residues (residues 69-78 of the spike protein) on the NTD calculated per chain. NIEA binding results in decreased epitope mobility. The red trace (chain A) represents the NIEA bound chains, with chain B in blue and chain C in green. Mean and 95% confidence interval are shown. Confidence interval was calculated using Seaborn ^36^.

### IgG binding results in stabilization of epitope residues

We observed a modest decrease in the mobility of the NIEA epitope residues on the NTD upon NIEA binding, as seen from the RMSF (Figure 4B). This is line with earlier MD simulations and HDX-MS experiments on the interactions between the SARS-CoV-2 spike NTD domain and the NIEA Fab^31^, which showed that NIEA binding stabilized a loop (loop 1) in the NTD^31^.

### Antibody cross-linked NTD-RBD pairs are de-coupled compared to unbound NTD-RBDs

There are three NTD – RBD pairs in a spike trimer, one of which (A-C) is NIEA bound. Each pair was monitored as a function of time by counting the number of interdomain contacts (defined as two atoms within a 0.3 nm range) and by measuring the distance between the geometric centers of the two domains. We observed a net increase in the NTD-RBD distance in the A-C pair (Figure 5). The NTD-RBD distances increased significantly (p=3×10^-8^) as a function of time for NIEA-bound NTD domains, compared to unbound domains (Figure 6A). Consistently, the number of contacts also decreased significantly (p= 1×10^-5^) as a function of time for antibody bound NTD domains, compared to unbound domains (Figure 7A). These observations were consistent across ten independent simulations. In most simulations there was an asymmetry between the spikes, such that NTD-RBD decoupling was stronger in one of the spikes, suggesting that decoupling of one NTD-RBD pair is sufficient to relax the system. In Figures 6-7, the NTD that was most decoupled from its neighboring RBD was selected and the non-bound NTD-RBD pairs were merged, resulting in the number of datapoints being double that of the bound NTDs in the calculation of the p-values. However, the differences in distance and numbers of contacts were statistically significant (p=2×10^-6^ and p=3×10^-5^ respectively) even when using the NTD-RBD pairs from both spikes (see supplemental Figures S2 & S3).

**Figure 5:**
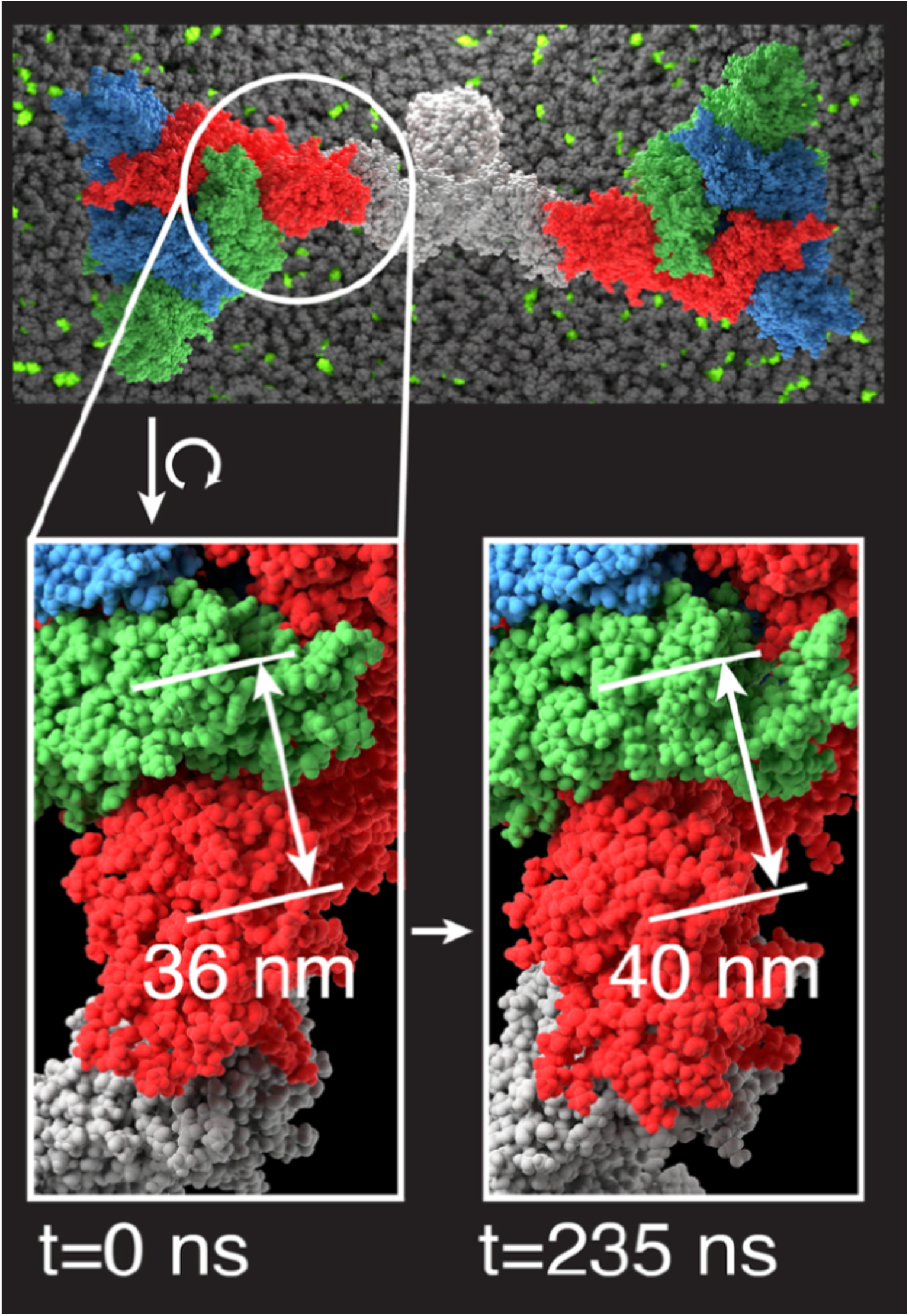
Increase in NTD-RBD distance during simulation. Top view of the IgG system, with insets zooming in on the NTD-RBD region of the IgG bound chain in one of the spikes. Shown are the NTD A (red), RBD C (green) and the IgG (grey). Bidirectional arrows indicate the distance between the center of geometry of the NTD and RBD domains. Glycans and solvent omitted for clarity.

**Figure 6:**
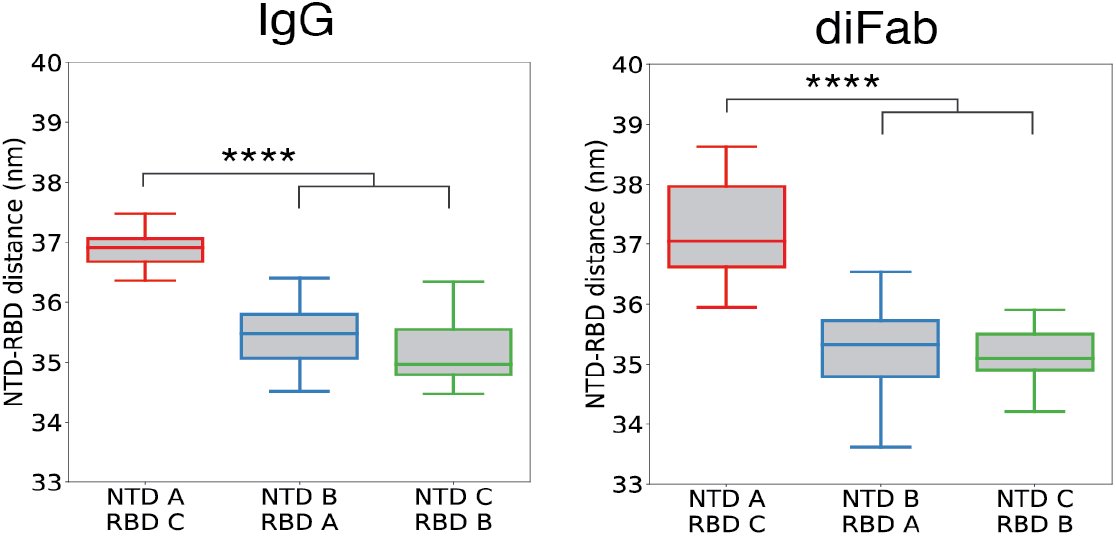
Distance between RBD-NTD in the IgG and diFab simulations. These boxes present the number of contacts for each of the two spikes over 10 independent simulations. There was in most simulations a strong asymmetry between the two spikes regarding the level of decoupling. To illustrate the effect of NIEA binding clearly, for boxes that represent the NIEA bound chain (red boxes), the spike with the strongest effect was selected. This means that the NIEA bound boxes only contain half of the datapoints with respect to the other two chains. Outliers are not shown. Please refer to supplemental Figure S2 for the boxplots which contain all datapoints.

**Figure 7:**
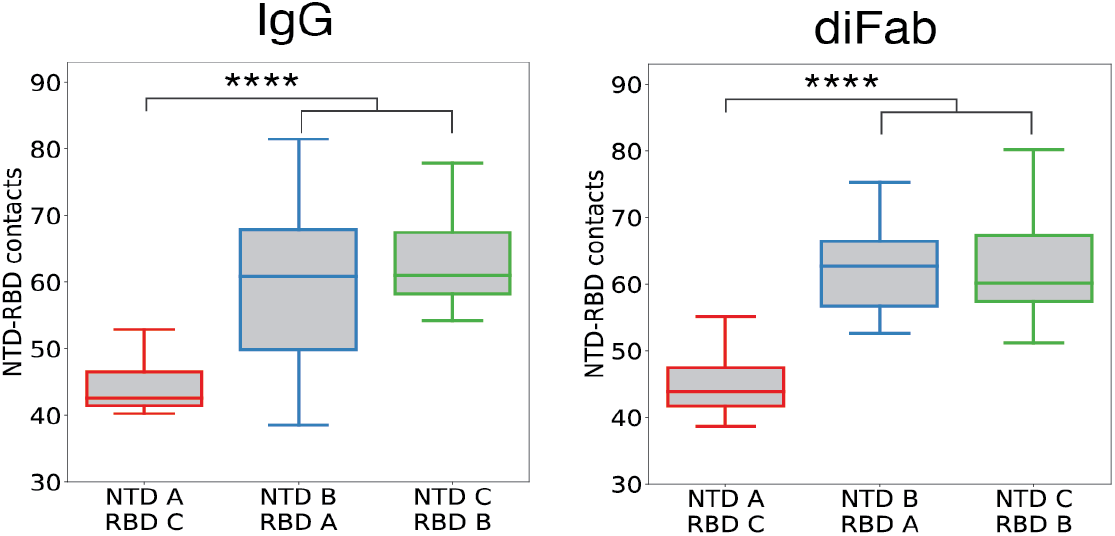
Number of contacts between RBD-NTD in the IgG and diFab simulations. These boxes present the number of contacts for each of the two spikes over 10 independent simulations. There was in most simulations a strong asymmetry between the two spikes regarding the level of decoupling. To illustrate the effect of NIEA binding clearly, for boxes that represent the NIEA bound chain (red boxes), the spike with the strongest effect was selected. This means that the NIEA bound boxes only contain half of the datapoints with respect to the other two chains. Atoms within a 0.3 nm radius were considered to be in contact. **** refers to a p-value < 0.0001. Outliers are not shown. Please refer to supplemental Figure S3 for the boxplots which contain all datapoints.

### NTD-RBD de-coupling is Fc independent

To study the effect of the Fc, we next examined the diFab system. The simulations of the diFab system were consistent with the observations above. There was a significant decrease in NTD-RBD contacts in the diFab bound chains (p=4×10^-7^) (Figure 7B) and increase in NTD-RBD separation (p=2×10^-11^) (Figure 6B). These results were consistent with previous experiments showing that the diFab is at least as effective as the IgG in enhancing ACE2 binding^13^. However, the Fc domain does influence the overall dynamics of the system. Specifically, the Fc was the most mobile part of the system, as evidenced by a higher RMSD for the IgG compared to the diFab (Figure 8), consistent with the previous EM results ^31^. As in the IgG simulations, there was little change in the RMSD of the spike proteins themselves upon diFab binding (supplemental Figure S4). In the simulations we frequently observed that the Fc domain would bend and or interact with the spike protein, which may also subtly influence the NTD-RBD separation.

**Figure 8:**
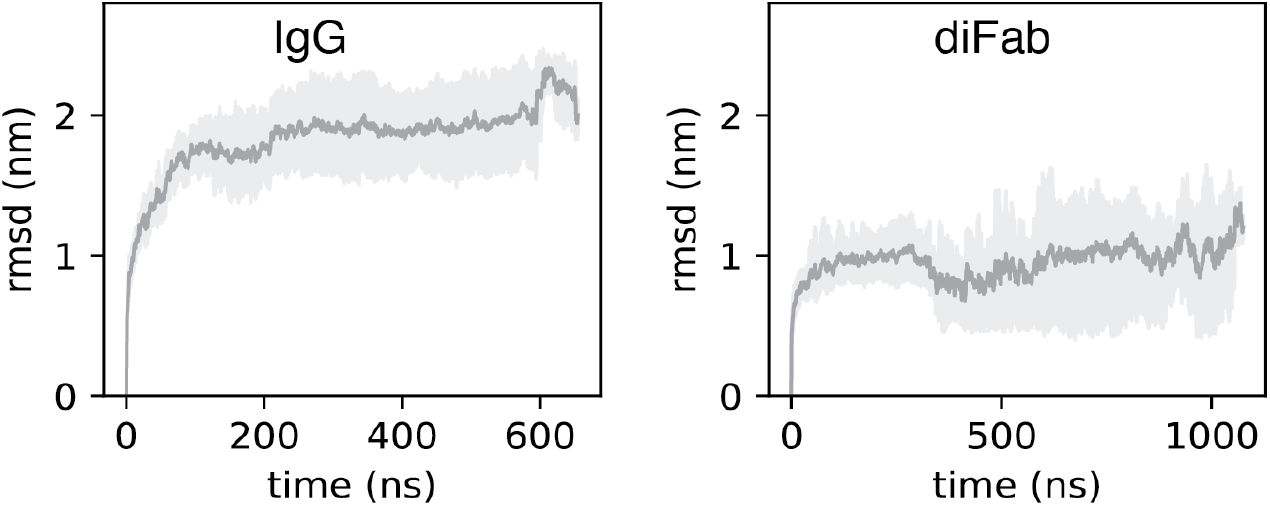
RMSD IgG and diFab. RMSD was calculated by fitting the trajectory on respectively the IgG and diFab. Mean and 95% confidence interval are shown.

## Discussion

Based on previous experimental work^31^, we extended our EM analysis of NIEA structure in order to understand full-length spike cross-linking. This resulted in the first direct observation of spike trimer cross-linking by negative staining EM imaging and demonstrated that addition of NIEAs to soluble spike trimers results in rosette formation. Moreover, we obtained a clear 2D class of a top-view of two cross-linked spikes using cryo-EM image reconstruction.

Taken together, the previous computational^13^, and domain based EM^31^ models are consistent with the cryo-EM analysis of full-length spikes and structural modeling here: NIEAs target an epitope in the NTD that conveniently allows cross-linking to neighboring spike trimers.

Based on this direct evidence, we next carried out extensive molecular dynamics simulations using a realistic model of NIEA-cross-linked spike trimers. Molecular dynamics studies have aided the understanding of the SARS-CoV-2 spike protein in various ways. Multiple molecular dynamics studies have demonstrated the important role of glycans in the infection process^11,34,37–39^. Weighted ensemble dynamics and targeted molecular dynamics simulations were used to characterize the down to up transition of the RBD domain^11,40^, and simulations of the complete SARS-CoV-2 envelope revealed how the different proteins interact in an intact virus^41–44^. Here, we constructed a minimal cross-linked system, consisting of two spike trimers joined by a single NIEA. The arrangement was assembled piecewise, with relaxation at each step, in order to minimize strain on the system. The simulations, which were carried out in realistic membrane and aqueous layers, were neither constrained nor driven toward a pre-established outcome. We paid particular attention to the NTD-RBD interactions, which might influence the RBD down/up transition^26–30^. Whether examined from the point of view of interdomain contacts or mean domain displacement, NTD-RBD interactions were significantly more decoupled in NIEA-bound NTDs than unbound NTDs, supporting our earlier hypothesis of the NIEA mode of action. By combining these observations with previous modeling, our results are consistent with a model in which NIEA binding facilitates RBD transition from a “down” to an “up” orientation.

Antibodies closely resembling NIEAs were previously reported by Barnes and co-workers; however, cross-linking was deemed unlikely based on modelling attempts using structural constraints^45^. However, this reasoning was based on an assumed spike-spike distance of 15 nm, which was subsequently found to be the *minimum* spacing between spikes in SARS-CoV^46,47^. More recent work has shown that mean spike-spike distance is closer to 23.6nm (StdDev=8.1)^32^, which agrees very well with the spike-spike distances established by our cross-linked models. Although the antibody COV57, as reported by Barnes et al., differs at the sequence level from known NIEAs, its geometry and binding angle are nearly identical (supplemental Figure S5). Given the fact that the 11 known NIEAs^12,13^ also vary significantly at the sequence level, along with close resemblance of their binding modes, we speculate that the NTD-targeting antibodies identified by Barnes and co-workers may indeed be NIEAs. Future work will clarify this question.

The spike-spike distance observed in our cryo-EM images was 13.9 nm, significantly smaller than the 21.7 nm used in the cross-linked system created by the modelling pipeline and the 23.6 nm measured by cryo-EM tomography^32^. We speculate that this discrepancy is due to the fact that the spike proteins in our experiments were not embedded in a membrane, allowing the entire system to relax further than when constrained by membrane embedding. Previously we showed that, as expected, the Fab arms of NTD-bound NIEAs are very flexible^31^. By changing the angle between the two Fab arms, an NIEA can cross-link spike proteins separated by a range of distances (Figure 9). The lack of membrane anchoring allows the entire system to minimize its surface area by minimizing spike trimer-trimer distance.

**Figure 9:**
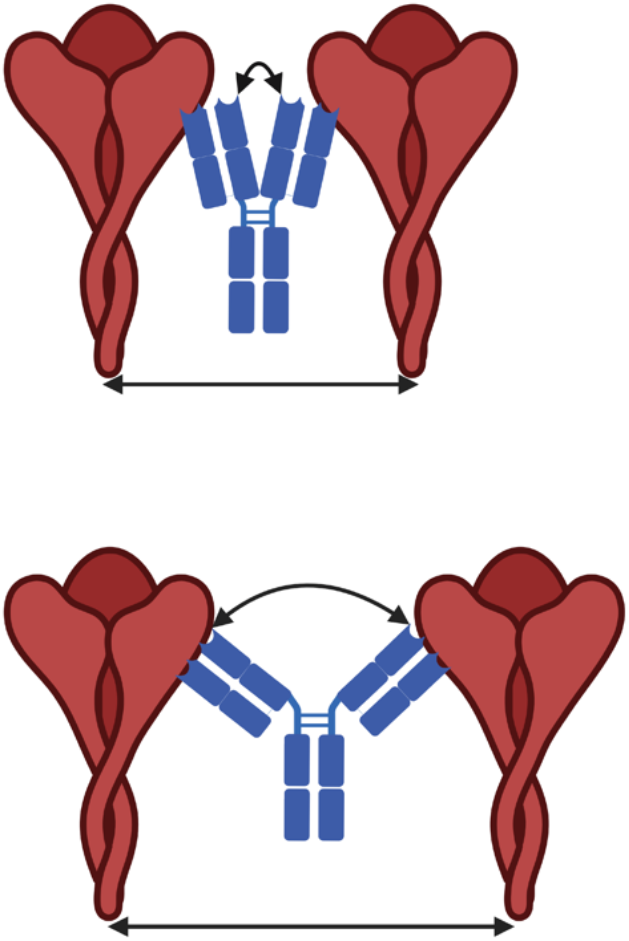
NIEAs can cross-link spike proteins separated by a range of distances. NIEAs can vary the angle between their Fab domains and can thereby cross-link both closely and further separated SARS-CoV-2 spike proteins. Schematic diagram was created with BioRender.com.

In this study a full 3D reconstruction of the NIEA cross-linking of two spike proteins was not feasible. Spike proteins notoriously adopt preferential orientations in cryo-EM grids^6,7^. The complex geometry of the system, combined with structural heterogeneity observed in multiple 2D classes, likely complicated the fitting and classification of a full high-resolution reconstruction. Nevertheless, by using selected 2D classes, it was possible to demonstrate the existence of spike-spike cross-linking.

While the role of ADE in SARS-CoV-2 *in vivo* is still controversial^12^, mutations can result in variants which are more susceptible to ADE^48–50^. Moreover, enhancing antibodies decrease the activity of neutralizing antibodies^13^. Although our research focused on SARS-CoV-2, ADE via cross-linking could also occur in other viruses for which the spike protein undergoes a similar conformational change upon host cell entry. Apart from coronaviruses like MERS-CoV and SARS-CoV^51^, influenza viruses^52–54^ undergo a conformational changes during host cell entry. It is therefore important in vaccine development to prevent the generation of infectivity enhancing antibodies. In the context of NIEA-mediated ADE, this might be as simple as mutating key NIEA epitope residues.

Taken together, this study provides the first direct structural and dynamic evidence for a novel Fc region-independent ADE mechanism. Given the fact that the frequency of NIEAs in COVID-19 patients is similar to that in vaccinated donors, but much less in healthy pre-pandemic donors^55^, the principles described in this investigation may inform future development of antibody-inducing vaccines.

## Methods

### Structural modelling of spike protein complex cross-linked by an infection-enhancing antibody

A trimeric spike model with glycosylation was adopted from the work of Casalino et al. (https://amarolab.ucsd.edu/files/covid19/PSF-PDB_spike_closed_prot_glyc_memb_wat_ions_amarolab.tar.gz)^34,40^. This structure was used as the main template for structure modelling. The PDB structure 7DZY was used as the template for the binding mode between a spike NTD and a Fab domain of 2490 antibody^13^, MODELLER^35^ was used to model to correct sequence. A random NTD domain and a multiple-template modelling pipeline of MODELLER^35^ was then applied to assemble NTD-Fab structure into the main template. In addition, missing residues within two other non-Fab binding NTDs were rebuilt by MODELLER. The grafted Fab-binding NTD structure lacks glycosylation, therefore missing glycan molecules from the main template were joined back into the model. Clash resulted from re-introduced glycan molecules were removed by resampling target residues’ sidechain orientation based on a rotamer library^33^. Based on the model of a trimeric spike in contact with Fab domain, a complex in which an antibody cross-links two spikes was built. Two spike/Fab models were positioned onto an artificial membrane plane by referring to their transmembrane domains. Two variables were used to position the two spikes so that they could be cross-linked by an enhancing antibody: 1) the relative rotational orientation of the two spikes, and 2) the distance between C-termini of the two Fab heavy chains. For each spike, principal coordinates were computed based on their central helix domain. Different rotation angles were sampled step wisely by rotating the spikes along the principal coordinate axis perpendicular to the plane. The distance between C-termini of the two Fabs was also sampled step wisely. For each combination of rotation and distance, a complete spike antibody model was generated by positioning an Fc domain in the vicinity of the Fab fragments and concatenating the Fc domain onto two Fab domains using MODELLER. The quality of each model was measured by calculating the RMSD of the two Fab domains upon introduction of the Fc domain. The model with the smallest RMSD was selected as the initial structure for the molecular dynamics simulations.

### Cell lines, plasmid, primers

Expi293 cells (Thermo Fisher Scientific) were cultured with the HE400AZ medium (Gmep). The cells were routinely checked for mycoplasma contamination. SARS-CoV-2 spike and 2490 monoclonal antibody plasmids were prepared as described previously^13^.

### Protein expression and purification

#### Trimer Spike

The pcDNA3.4 plasmid containing His-tagged SARS-CoV-2 spike protein was transfected to Expi293T cells using PEI max (Polysciences). After 18–21 h post-transfection, Gxpress 293 Enhancer (Gmep) was added into culture media to enhance translation. Culture supernatant containing His-tagged NTD protein was harvested 4 days after transfection and was purified using Ni-NTA resin (Qiagen). Spike protein was further purified with a Superose 6 Increase 10/300 GL column equilibrated with 20 mM Tris-HCl (pH 8.0), 300 mM NaCl using AKTA Pure 25 System (Cytiva). Fraction under the gel filtration peak was concentrated using Ultrafiltration unit MWCO 100 (Sartorius).

#### 2490 Antibody

The pCAGGS vector containing the 2490 antibody was transfected using PEI into Expi293T cells. Gexpress 293 (Gmep) enhancer was added into culture media after 18–21 h post-transfection. Culture supernatant containing the 2490 antibody was harvested 4 days post-transfection and purified with HiTrap Protein A column using AKTA Pure 25 System (Cytiva) followed by buffer exchange into PBS.

#### 2490 antibody - spike trimer complex formation and purification

The spike trimer was incubated with the 2490 antibody at a molar ratio of 1:1 or 2:1and incubated at room temperature for 16–18 h. The complex was purified on a Superdex 75 Increase 10/300 GL (Cytiva) column equilibrated with 20 mM Tris-HCl pH 8.0, 300 mM NaCl. The purified complex was used for negative staining electron microscopy or cryo-EM analysis.

### Negative Staining Electron Microscopy

An aliquot of 5 μL of purified complex was applied to a glow discharged, 600-mesh, carbon-coated grid for 30 s. The grid was then blotted with filter paper to remove excess sample. Subsequently, the grid was negatively stained with three drops of 2% uranyl acetate, blotted again with filter paper, and air-dried. Finally, the grid was transferred to a JEOL 1400 plus transmission electron microscope equipped with a Gatan OneView CMOS camera and operated at 80 kV.

### CryoEM imaging and data analysis

Cryo-grid was glow discharged at 9 mA for 30 seconds. 2.5 μl complex of spike and antibody (0.17 mg/ml) solution was applied onto the Quantifoil Au 0.6/1.0 200 mesh cryo-grid and frozen in liquid ethane using a Vitrobot IV (FEI, 6°C and 95% humidity). Data collection of each sample was carried out on a Titan Krios (FEI, Netherlands) equipped with a thermal field emission electron gun operated at 300 kV, an energy filter with a 20 eV slit width and a K3 direct electron detector camera (Gatan, USA). For automated data acquisition, SerialEM software was used to collect cryo-EM image data. Movie frames were recorded using the K3 camera at a calibrated magnification of × 81,000 corresponding to a pixel size of 0.88 Å with a setting defocus range from -0.6 to -2.0 μm. The data were collected with a total exposure of 3 s fractionated into 50 frames, with a total dose of ∼50 electrons Å^2^ in counting mode. A total of 10,898 movies of NIEA—Spike protein complex were collected.

### Image processing

All image processing was carried out using cryoSPARC software v4.3.0^56^. After motion correction, CTF estimation, and manual curation, 914,329 particles were automatically picked using Topaz picking algorithm. The particles were subjected to 2D classified into 200 classes, then 801,093 particles were selected after 2D classification. After heterogeneous refinement into 6 classes using Ab-initio reconstructed map as reference maps, 2 density maps for spike-NIEA complex were obtained. In one of the density maps, Fab from NIEA bound to one of trimeric spike protomers. In another map, all of three protomers occupied by Fabs from NIEA. Since we know that there are particles of spike protein complexed with NIEA, to focus the complex of spike-NIEA-spike, we performed re-extraction of particles by shifting center of particle box closer to center of IgG density. The shifting values were measured by Makers tool using UCSF ChimeraX^57^. Then the particle coordinates were shifted using Volume Alignment Tools in cryoSPARC. Following particle re-extraction, several rounds of 2D classification was carried out to select good classes.

### Molecular dynamics

The topologies provided by Casalino et al proved to be incompatible with Gromacs and therefore were re-created using Charmm-gui^58,59^. The spike proteins were glycosylated using the glycan reader & modeler tools of Charmm-gui^60,61^, according to the glycosylation pattern described by Casalino et al.^34^. The obtained parametrized spikes and antibody were fitted upon the system provided by Li et al, and subsequently the whole system was embedded in a membrane with the composition according to Casalino et al.^34^, see Table 2. Both membrane leaflets were composed out of 1800 lipids. For the diFab the residues up to Cys 236 from IgG 2490 were used. The IgG system was solvated in a solution of 1020739 water molecules and 3449 NaCl molecules, corresponding to ∼0.154 M, which is a physiological saline solution, an additional 720 Na^+^ counterions were added to neutralize charges. The diFab system was solvated in 1022974 and 3449 NaCL ions and an additional 720 Na^+^ counterions. Simulations were performed with the Charmm36m forcefield (February 2021 release)^62^. Equations of motions were integrated with Gromacs 2022.x ^63,64^. Simulation parameters were as provided by Charmm-gui, with the modification that simulations only proved to be stable with a timestep of 1 fs, and that velocities were generated at the start of each run. In short: the Verlet cutoff scheme was used with a cutoff of 1.2 nm and the neighbor list updated every 20 steps. Van der Waals interactions were calculated up to 1.2 nm using a plain cut-off, but with the forces being switched to zero in the 1 to 1.2 nm range. Short range electrostatics were calculated up to a 1.2 nm cut-off and Particle-Mesh Ewald was used for the long range electrostatics^65^. Pressure was kept semiistropically at p= 1.0 bar using the Parrinello-Rahman barostat^66^ with a τ_p_ = 5.0 ps and a compressibility of 4.5 ×10^-5^ bar^-1^. Thermostat was kept at 310 K using the Nosé-Hoover thermostat^67,68^ with τ_t_ = 1.0 ps and protein, membrane and solvent coupled separately. Bonds involving hydrogen atoms were constrained using the LINCS algorithm^69^. Periodic boundary jumps were corrected by using a combination of MDwhole^70^ and the Gromacs tool gmx trjconv.

**Table 1:**
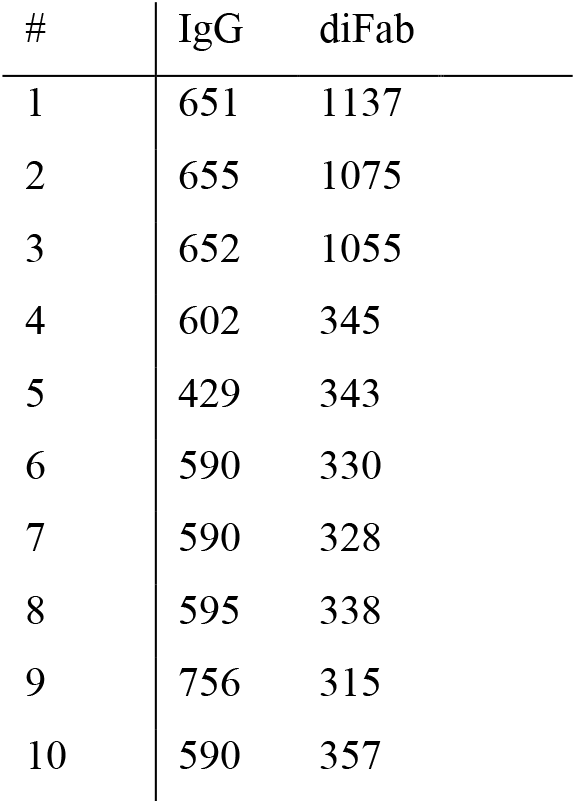
Simulation length in nanoseconds for the 10 independent IgG and diFab simulations.

**Table 2:** Lipid composition of the inner and outer leaflet of the IgG and diFab simulation systems.

| # | Inner leaflet | Outer leaflet |
| --- | --- | --- |
| CHL1 | 270 | 270 |
| POPC | 846 | 846 |
| POPE | 360 | 360 |
| POPI | 198 | 198 |
| POPS | 126 | 126 |

### Analysis

RMSD was calculated over spike and IgG/diFab backbone atoms after fitting the trajectories on the protein backbone. RMSF was calculated for the Cα atoms of the epitope residues HVSGTNGTKR. Contacts were calculated between all atoms, atoms within a 0.3 nm cutoff were considered to be in contact. Distances were calculated between all atoms. Analysis was performed with the Gromacs package,MDAnalysis^71,72^ and Seaborn^36^.

Figures of molecular dynamics simulations were rendered with Blender^73^ using the Molecular Nodes plugin^74^.

### Sequence analysis and spike-Fab binding angle comparison

Sequence analysis was performed using the Geneious Prime software (version 2024.0.7). Multiple alignments were done using MAFFT^75^. Sequences of NIEAs were taken from previous studies ^13, 12^ and sequence of COV57 antibody was taken from article published by Rabbani et al.^76^.

Structures of trimeric spike-Fab complex from previous studies ^12,13,45^were retrieved from RCSB PDB (PDB ID: 7LAB, 7LCN, 7DZY, 7DZX) or from EMDB (EMDB ID: <u>EMD-</u> <u>22936</u>, <u>EMD-22942</u>, <u>EMD-22125</u>).

## Acknowledgements

This work used computational resources of the supercomputer Fugaku provided by RIKEN through the HPCI System Research Project (Project ID: hp220148) and the TSUBAME3.0 supercomputer at Tokyo Institute of Technology. HA and DMA acknowledge funding from CAMaD, grant number JP223fa627002.

## Bibliography

1. Massey, D., Berrent, D. & Krumholz, H. Breakthrough Symptomatic COVID-19 Infections Leading to Long Covid: Report from Long Covid Facebook Group Poll. medRxiv (2021).

2. Tan, S. T. et al. Infectiousness of SARS-CoV-2 breakthrough infections and reinfections during the Omicron wave. Nat Med 29, 358–365 (2023).

3. Al-Aly, Z., Bowe, B. & Xie, Y. Long COVID after breakthrough SARS-CoV-2 infection. Nat Med 28, 1461–1467 (2022).

4. Khoury, D. S. et al. Neutralizing antibody levels are highly predictive of immune protection from symptomatic SARS-CoV-2 infection. Nat Med 27, 1205–1211 (2021).

5. Garcia-Beltran, W. F. et al. COVID-19-neutralizing antibodies predict disease severity and survival. Cell 184, 476–488.e11 (2021).

6. Wrapp, D. et al. Cryo-EM structure of the 2019-nCoV spike in the prefusion conformation. Science 367, 1260–1263 (2020).

7. Shang, J. et al. Cell entry mechanisms of SARS-CoV-2. Proc Natl Acad Sci U S A 117, 11727–11734 (2020).

8. Henderson, R. et al. Controlling the SARS-CoV-2 spike glycoprotein conformation. Nat Struct Mol Biol 27, 925–933 (2020).

9. Wrapp, D. et al. Structural Basis for Potent Neutralization of Betacoronaviruses by Single-Domain Camelid Antibodies. Cell 181, 1004–1015.e15 (2020).

10. Lu, M. et al. Real-Time Conformational Dynamics of SARS-CoV-2 Spikes on Virus Particles. Cell Host Microbe 28, 880–891.e8 (2020).

11. Mori, T. et al. Elucidation of interactions regulating conformational stability and dynamics of SARS-CoV-2 S-protein. Biophys J 120, 1060–1071 (2021).

12. Li, D. et al. In vitro and in vivo functions of SARS-CoV-2 infection-enhancing and neutralizing antibodies. Cell (2021).

13. Liu, Y. et al. An infectivity-enhancing site on the SARS-CoV-2 spike protein targeted by antibodies. Cell 184, 3452–3466.e18 (2021).

14. Bournazos, S., Gupta, A. & Ravetch, J. V. The role of IgG Fc receptors in antibody-dependent enhancement. Nat Rev Immunol 20, 633–643 (2020).

15. Wang, T. T. et al. IgG antibodies to dengue enhanced for FcγRIIIA binding determine disease severity. Science 355, 395–398 (2017).

16. Vennema, H. et al. Early death after feline infectious peritonitis virus challenge due to recombinant vaccinia virus immunization. Journal of virology 64, 1407–1409 (1990).

17. Weiss, R. C. & Scott, F. W. Antibody-mediated enhancement of disease in feline infectious peritonitis: comparisons with dengue hemorrhagic fever. Comp Immunol Microbiol Infect Dis 4, 175–189 (1981).

18. Iwasaki, A. & Yang, Y. The potential danger of suboptimal antibody responses in COVID-19. Nat Rev Immunol 20, 339–341 (2020).

19. Arvin, A. M. et al. A perspective on potential antibody-dependent enhancement of SARS-CoV-2. Nature 584, 353–363 (2020).

20. Haynes, B. F. et al. Prospects for a safe COVID-19 vaccine. Science Translational Medicine 12, eabe0948 (2020).

21. Nakayama, E. E. & Shioda, T. SARS-CoV-2 Related Antibody-Dependent Enhancement Phenomena In Vitro and In Vivo. Microorganisms 11, 1015 (2023).

22. Duncan, A. R. & Winter, G. The binding site for C1q on IgG. Nature 332, 738–740 (1988).

23. Okuya, K. et al. Multiple Routes of Antibody-Dependent Enhancement of SARS-CoV-2 Infection. Microbiol Spectr 10, e0155321 (2022).

24. Wan, Y. et al. Molecular Mechanism for Antibody-Dependent Enhancement of Coronavirus Entry. J Virol 94, e02015–19 (2020).

25. Ziganshina, M. M. et al. Antibody-Dependent Enhancement with a Focus on SARS-CoV-2 and Anti-Glycan Antibodies. Viruses 15, 1584 (2023).

26. Qing, E. et al. Dynamics of SARS-CoV-2 Spike Proteins in Cell Entry: Control Elements in the Amino-Terminal Domains. mBio 12, e0159021 (2021).

27. Gobeil, S. M. et al. Effect of natural mutations of SARS-CoV-2 on spike structure, conformation, and antigenicity. Science 373, eabi6226 (2021).

28. Braet, S. M. et al. Timeline of changes in spike conformational dynamics in emergent SARS-CoV-2 variants reveal progressive stabilization of trimer stalk with altered NTD dynamics. Elife 12, e82584 (2023).

29. Fallon, L. et al. Free Energy Landscapes from SARS-CoV-2 Spike Glycoprotein Simulations Suggest that RBD Opening Can Be Modulated via Interactions in an Allosteric Pocket. J Am Chem Soc 143, 11349–11360 (2021).

30. Ray, D., Le, L. & Andricioaei, I. Distant residues modulate conformational opening in SARS-CoV-2 spike protein. Proc Natl Acad Sci U S A 118, e2100943118 (2021).

31. Lusiany, T. et al. Enhancement of SARS-CoV-2 Infection via Crosslinking of Adjacent Spike Proteins by N-Terminal Domain-Targeting Antibodies. Viruses 15, 2421 (2023).

32. Klein, S. et al. SARS-CoV-2 structure and replication characterized by in situ cryo-electron tomography. Nat Commun 11, 5885 (2020).

33. Hintze, B. J., Lewis, S. M., Richardson, J. S. & Richardson, D. C. Molprobity’s ultimate rotamer-library distributions for model validation. Proteins 84, 1177–1189 (2016).

34. Casalino, L. et al. Beyond Shielding: The Roles of Glycans in the SARS-CoV-2 Spike Protein. ACS Cent Sci 6, 1722–1734 (2020).

35. Webb, B. & Sali, A. Comparative Protein Structure Modeling Using MODELLER. Curr Protoc Bioinformatics 54, 5.6.1–5.6.37 (2016).

36. Waskom, M. seaborn: statistical data visualization. Journal of Open Source Software 6, 3021 (2021).

37. Turoňová, B. et al. In situ structural analysis of SARS-CoV-2 spike reveals flexibility mediated by three hinges. Science 370, 203–208 (2020).

38. Choi, Y. K. et al. Structure, Dynamics, Receptor Binding, and Antibody Binding of the Fully Glycosylated Full-Length SARS-CoV-2 Spike Protein in a Viral Membrane. J Chem Theory Comput 17, 2479–2487 (2021).

39. Harbison, A. M. et al. Fine-tuning the spike: role of the nature and topology of the glycan shield in the structure and dynamics of the SARS-CoV-2 S. Chem Sci 13, 386–395 (2022).

40. Sztain, T. et al. A glycan gate controls opening of the SARS-CoV-2 spike protein. Nat Chem 13, 963–968 (2021).

41. Yu, A. et al. A multiscale coarse-grained model of the SARS-CoV-2 virion. Biophys J 120, 1097–1104 (2021).

42. Wang, B., Zhong, C. & Tieleman, D. P. Supramolecular Organization of SARS-CoV and SARS-CoV-2 Virions Revealed by Coarse-Grained Models of Intact Virus Envelopes. J Chem Inf Model 62, 176–186 (2022).

43. Wang, D. et al. Toward Atomistic Models of Intact SARS-CoV-2 via Martini Coarse-Grained Molecular Dynamics Simulations. bioRxiv (2022).

44. Pezeshkian, W. et al. Molecular architecture and dynamics of SARS-CoV-2 envelope by integrative modeling. Structure 31, 492–503.e7 (2023).

45. Barnes, C. O. et al. Structures of Human Antibodies Bound to SARS-CoV-2 Spike Reveal Common Epitopes and Recurrent Features of Antibodies. Cell 182, 828–842.e16 (2020).

46. Neuman, B. W. et al. Supramolecular architecture of severe acute respiratory syndrome coronavirus revealed by electron cryomicroscopy. J Virol 80, 7918–7928 (2006).

47. Neuman, B. W. et al. A structural analysis of M protein in coronavirus assembly and morphology. J Struct Biol 174, 11–22 (2011).

48. Kimura, I. et al. The SARS-CoV-2 Lambda variant exhibits enhanced infectivity and immune resistance. Cell Rep 38, 110218 (2022).

49. Liu, Y. et al. The SARS-CoV-2 Delta variant is poised to acquire complete resistance to wild-type spike vaccines. bioRxiv (2021).

50. Liu, Y. & Arase, H. Neutralizing and enhancing antibodies against SARS-CoV-2. Inflamm Regen 42, 58 (2022).

51. Yuan, Y. et al. Cryo-EM structures of MERS-CoV and SARS-CoV spike glycoproteins reveal the dynamic receptor binding domains. Nat Commun 8, 15092 (2017).

52. Das, D. K. et al. Direct Visualization of the Conformational Dynamics of Single Influenza Hemagglutinin Trimers. Cell 174, 926–937.e12 (2018).

53. Garcia, N. K. et al. Structural dynamics reveal subtype-specific activation and inhibition of influenza virus hemagglutinin. J Biol Chem 299, 104765 (2023).

54. Dam, K. A., Fan, C., Yang, Z. & Bjorkman, P. J. Intermediate conformations of CD4-bound HIV-1 Env heterotrimers. Nature 623, 1017–1025 (2023).

55. Ismanto, H. S. et al. Landscape of infection enhancing antibodies in COVID-19 and healthy donors. Comput Struct Biotechnol J 20, 6033–6040 (2022).

56. Punjani, A., Rubinstein, J. L., Fleet, D. J. & Brubaker, M. A. cryoSPARC: algorithms for rapid unsupervised cryo-EM structure determination. Nature Methods 14, 290–296 (2017).

57. Pettersen, E. F. et al. UCSF ChimeraX: Structure visualization for researchers, educators, and developers. Protein Sci 30, 70–82 (2021).

58. Jo, S., Kim, T., Iyer, V. G. & Im, W. CHARMM-GUI: a web-based graphical user interface for CHARMM. Journal of computational chemistry 29, 1859–1865 (2008).

59. Lee, J. et al. CHARMM-GUI input generator for NAMD, GROMACS, AMBER, OpenMM, and CHARMM/OpenMM simulations using the CHARMM36 additive force field. Journal of chemical theory and computation 12, 405–413 (2016).

60. Jo, S., Song, K. C., Desaire, H., MacKerell Jr, A. D. & Im, W. Glycan Reader: automated sugar identification and simulation preparation for carbohydrates and glycoproteins. Journal of computational chemistry 32, 3135–3141 (2011).

61. Park, S.-J. et al. CHARMM-GUI Glycan Modeler for modeling and simulation of carbohydrates and glycoconjugates. Glycobiology 29, 320–331 (2019).

62. Huang, J. et al. CHARMM36m: an improved force field for folded and intrinsically disordered proteins. Nat Methods 14, 71–73 (2017).

63. Hess, B., Kutzner, C., van der Spoel, D. & Lindahl, E. GROMACS 4: Algorithms for Highly Efficient, Load-Balanced, and Scalable Molecular Simulation. Journal of Chemical Theory and Computation 4, 435–447 (2008).

64. Abraham, M. J. et al. GROMACS: High performance molecular simulations through multi-level parallelism from laptops to supercomputers. SoftwareX 1, 19–25 (2015).

65. Darden, T., York, D. & Pedersen, L. Particle mesh Ewald: An N⋅ log (N) method for Ewald sums in large systems. The Journal of Chemical Physics 98, 10089–10092 (1993).

66. Parrinello, M. & Rahman, A. Polymorphic transitions in single crystals: A new molecular dynamics method. Journal of Applied Physics 52, 7182–7190 (1981).

67. Hoover, W. G. Canonical dynamics: Equilibrium phase-space distributions. Physical review A 31, 1695 (1985).

68. Nosé, S. A molecular dynamics method for simulations in the canonical ensemble. Molecular Physics 52, 255–268 (1984).

69. Hess, B., Bekker, H., Berendsen, H. J. C. & Fraaije, J. G. E. M. LINCS: a linear constraint solver for molecular simulations. Journal of computational chemistry 18, 1463–1472 (1997).

70. Bruininks, B. M. H., Wassenaar, T. A. & Vattulainen, I. Unbreaking Assemblies in Molecular Simulations with Periodic Boundaries. J Chem Inf Model (2023).

71. Gowers, R. et al. MDAnalysis: A Python Package for the Rapid Analysis of Molecular Dynamics Simulations. Proceedings of the Python in Science Conference 98–105 (2016).

72. Michaud-Agrawal, N., Denning, E. J., Woolf, T. B. & Beckstein, O. MDAnalysis: a toolkit for the analysis of molecular dynamics simulations. Journal of Computational Chemistry 32, 2319–2327 (2011).

73. Blender, O. C. Blender - a 3D modelling and rendering package (Blender Foundation, 2018).

74. Brady, J. et al. MolecularNodes: v2.7.4 for Blender 3.5+ (Zenodo, 2023).

75. Nakamura, T., Yamada, K. D., Tomii, K. & Katoh, K. Parallelization of MAFFT for large-scale multiple sequence alignments. Bioinformatics 34, 2490–2492 (2018).

76. Robbiani, D. F. et al. Convergent antibody responses to SARS-CoV-2 in convalescent individuals. Nature 584, 437–442 (2020).

