## Supplementary information for "Fc-independent SARS-CoV-2 infection-enhancing antibodies decouple N-terminal and receptor-binding domains by cross-linking neighboring spikes"

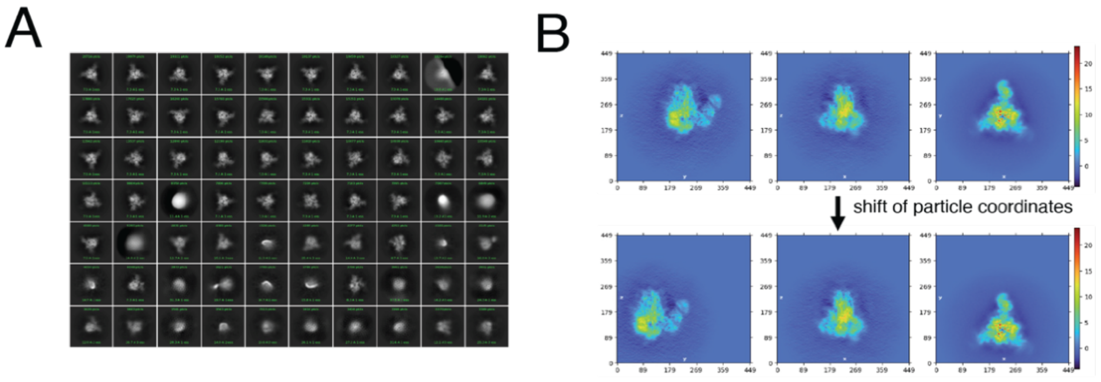


**Figure S1 cryo-EM images processing.** (A) 2D classes of trimeric spike protein complexed with NIEA before particle re-extraction. The density corresponds to trimeric spike protein and the Fab region of the NIEA IgG. (B) The center of the particle box was shifted from the center of the trimeric spike particles to the center of the IgG particles in order to re-extract particles.

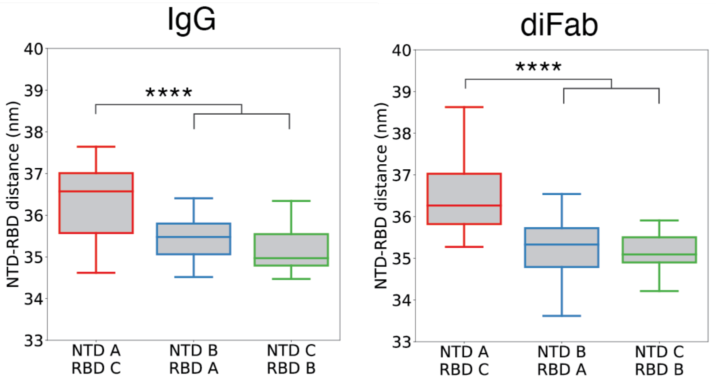


**Figure S2: Distance between RBD-NTD in the IgG and diFab simulations using the Ig bound NTD from both spikes.** These boxes present the number of contacts for each of the two spikes over 10 independent simulations. When using the Ig bound chains of both spikes in the p-value calculation, $2\times10^{-6}$ for the IgG and $2\times10^{-8}$ for the diFab.**** refers to a p-value < 0.0001.

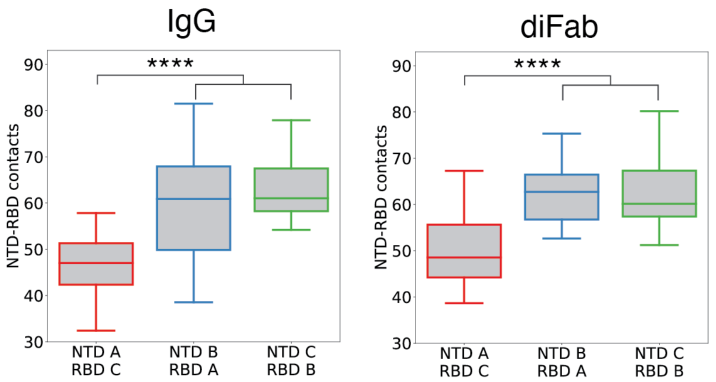

**Figure S3: Number of contacts between RBD-NTD in the IgG and diFab simulations using the Ig bound NTD from both spikes.** These boxes present the number of contacts for each of the two spikes over 10 independent simulations. When using the Ig bound chains of both spikes in the p-value calculation, $3\times10^{-5}$ for the IgG and $1\times10^{-5}$ for the diFab. Atoms within a 0.3 nm radius were considered to be in contact. **** refers to a p-value < 0.0001.

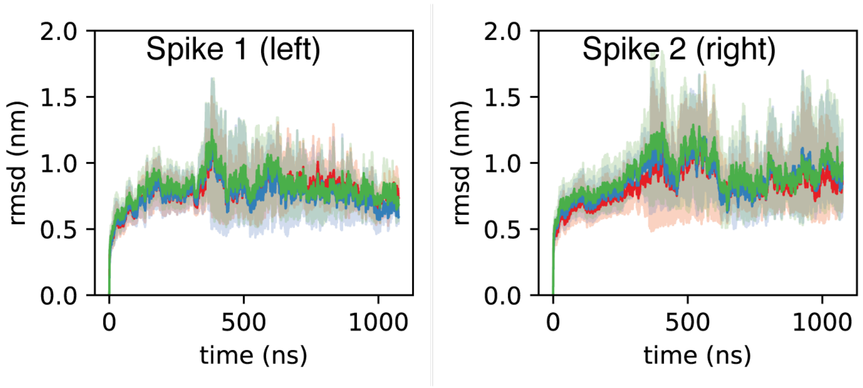


**Figure S4: RMSD of the SARS-CoV-2 spikes in the diFab bound simulation.** The red trace represents the diFab bound chains. RMSD was calculated by fitting the trajectory on the whole spike protein and the RMSD was calculated per chain. Mean and 95% confidence interval are shown. Please note that the spread in the confidence interval sharply increases around 350 ns, this is because only three IgG simulations were longer than 360 ns. Spike 1 (left) corresponds to the left spike in Figure 3A, and Spike 2 to the right one. Chain A (red), chain B (blue) and chain C (green).

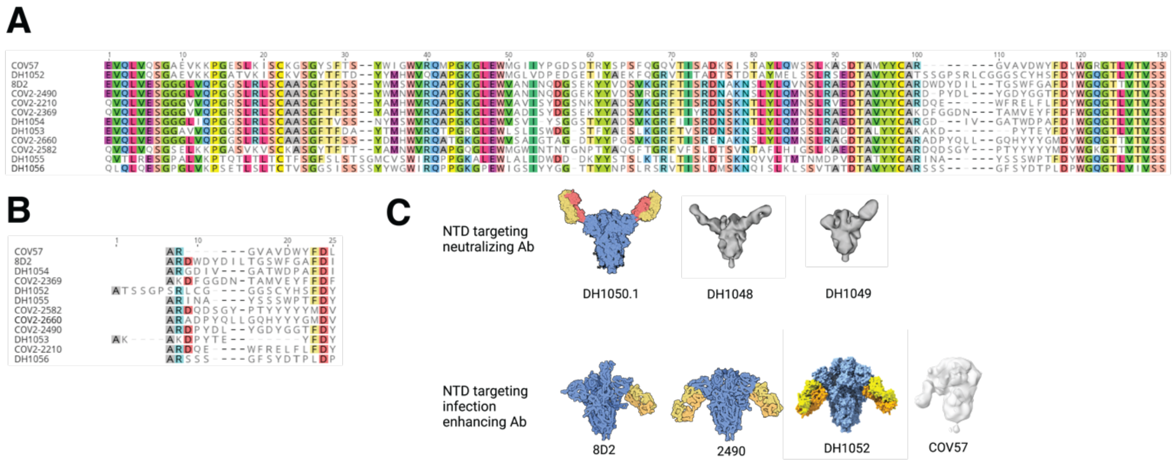


**Figure S5: Comparison of COV57 with known NIEAs.** (A) Multiple Sequence Alignment (MSA) of heavy chain variable (VH) region from COV57 and known NIEAs. (B) MSA of CDR3 region from COV57 and known NIEAs. (C) Binding angles between known NTD targeting neutralizing Ab, known NIEAs, and COV57.

Spike protein is colored in blue while Fab is colored in yellow, dark yellow, or orange. Density map of spike protein complexed with Fab are shown in grey or light grey. Structures of trimeric spike-Fab complex [1–3]were retrieved from RCSB PDB (PDB ID: 7LAB, 7LCN, 7DZY, 7DZX) or from EMDB (EMDB ID: EMD-22936, EMD-22942, EMD-22125). Schematic diagram was created with BioRender.com

### Supplemental Bibliography


1. Barnes, C. O. *et al.* Structures of Human Antibodies Bound to SARS-CoV-2 Spike Reveal Common Epitopes and Recurrent Features of Antibodies. *Cell* **182**, 828–842.e16 (2020).
2. Li, D. *et al.* In vitro and in vivo functions of SARS-CoV-2 infection-enhancing and neutralizing antibodies. *Cell* (2021).
3. Liu, Y. *et al.* An infectivity-enhancing site on the SARS-CoV-2 spike protein targeted by antibodies. *Cell* **184**, 3452–3466.e18 (2021).
